# Sulfasalazine promotes myogenic differentiation via increasing of fast myosin heavy chain isoforms

**DOI:** 10.1101/2023.08.15.553466

**Authors:** Seungju Cho, Seonggyu Choi, Hyunjun Chang, Nguyen Khoi Tran, Hyunji Lee, Sunyoung Ryu, Mihee Park, Sangchul Choi, Jisoo Park, Jongsun Park

## Abstract

Sarcopenia is a debilitating condition characterized by the progressive and generalized degeneration of skeletal muscle mass and function. As of now, there is no approved pharmacological treatment for sarcopenia. Previously, our research revealed that Yin Yang 1 (YY1) plays a crucial role in PHD finger protein 20 (PHF20)-mediated myogenic differentiation. A significant enhancement in YY1 transcription, mediated by PHF20, was observed in C_2_C_12_ myoblasts. Within skeletal muscle, YY1 is traditionally considered to inhibit myogenesis by directly repressing the synthesis of late-stage differentiation genes, such as skeletal muscle actin, muscle creatine kinase, and myosin heavy chain IIb. Through screening of a drug library using a PHF20/YY1 promoter reporter assay, sulfasalazine emerged as a promising candidate. Sulfasalazine is known for its anti-inflammatory and immunomodulatory properties and is commonly prescribed for autoimmune diseases. In this study, the treatment of C_2_C_12_ myoblasts with sulfasalazine accelerated the myogenic differentiation and bolstered the gene and protein expression of fast myosin heavy chain via a metabolic shift towards glycolysis. Additionally, oral administration of sulfasalazine demonstrated improvement in physical performance in aged mice, as well as in mice models with hindlimb disuse or damage. Moreover, sulfasalazine exhibited a remarkable ability to facilitate the recovery of muscle fibers damaged by Velcro immobilization. Collectively, our findings suggest that sulfasalazine could represent a novel therapeutic avenue for the amelioration of muscle weakness, including sarcopenia.

## Introduction

Sarcopenia is a progressive condition characterized by a generalized decline in skeletal muscle mass and function, leading to increased risks of disability, metabolic dysfunction, poor quality of life, and mortality [1]. Although sarcopenia is primarily associated with aging, various other factors including sedentary lifestyle, immobilization, malnutrition, diabetes, obesity, and acute or chronic inflammatory diseases contribute to muscle mass reduction [2], which in turn can exacerbate these conditions [3, 4]. Despite the severity of sarcopenia, the Food and Drug Administration (FDA) has not yet approved any specific drug for its treatment. Several agents such as growth hormone, anabolic or androgenic steroids, selective androgen receptor modulators, protein anabolic agents, appetite stimulants, myostatin inhibitors, Type II receptor activators, β-receptor blockers, angiotensin-converting enzyme inhibitors, and troponin activators are under consideration.

Sulfasalazine, a traditional medication for rheumatoid arthritis, has been widely administered to treat autoimmune diseases including ankylosing spondylitis, Crohn’s disease, and ulcerative colitis [5–8]. Although the exact mechanism remains unclear, sulfasalazine is known to exert anti-inflammatory and immunomodulatory effects [5].

YY1 is a DNA-binding protein with four C-terminal zinc finger domains, acting as either an activator or repressor of gene expression [9]. In skeletal muscle, rapamycin treatment, which inhibits the mammalian target of rapamycin (mTOR), led to reduced gene expression of mitochondrial transcriptional regulators such as peroxisome proliferator-activated receptor γ coactivator 1α (PGC-1α), estrogen-related receptor α, and nuclear respiratory factors, resulting in diminished mitochondrial gene expression and oxygen consumption [10]. YY1 is postulated to hinder myogenesis by directly repressing late-stage differentiation genes, including skeletal actin [11], muscle creatine kinase (MCK), and myosin heavy chain (MyHC) type IIb [12, 13]. Recent evidence suggests that YY1 binding and subsequent recruitment of the Polycomb suppressor complex, containing the Ezh2 methyl-transferase along with HDAC1, regulate the transcriptional silencing of MCK enhancer and MyHC type IIb promoter in proliferating myoblasts [12]. During skeletal myogenesis, the YY1/Ezh2/HDAC repressive complex is replaced by activators such as serum response factor and MyoD, and associated acetyl-transferases CBP and p300/CBP-associated factor (PCAF) on MCK and MyHC DNA. These activators are essential for the expression of late-stage differentiation genes. Our prior findings indicate that YY1 is essential for PHF20-mediated myogenic differentiation, with PHF20 promoting YY1 transcription in C_2_C_12_ myoblasts [14]. Moreover, it has been reported that YY1-binding motifs are highly enriched in mitochondrial genes regulated by rapamycin and PGC1α, and knockdown of YY1 resulted in decreased expression of PGC1α, mitochondrial genes, and oxygen consumption, suggesting PGC1α regulation by YY1 [10].

Skeletal muscle is a multifaceted tissue comprised of diverse fibers, each being a multinucleated syncytium originating from myoblast differentiation and fiber cell fusion. These fibers are classified into four primary types: MyHC types I, IIa, IIx, and IIb [15, 16], and are adapted to different bioenergetic and biophysical properties to meet varying functional demands during postnatal muscle growth and development. For instance, resistance training notably reduces mitochondrial content, promotes slow-to-fast fiber transformation, and shifts the energy production system towards glycolysis [17]. MyHC type I fibers are highly oxidative and contain abundant mitochondria, enabling sustained contractions with reduced fatigue. They favor fatty acid oxidation, a more efficient ATP production pathway compared to anaerobic glycolysis [18]. These fibers, rich in myoglobin and cytochromes, are predominantly found in deep portions of muscles such as the soleus and deep gastrocnemius (GA), making them well-suited for the slow and steady requirements of locomotion [19, 20]. By contrast, type IIb fibers, located at the other end of the metabolic spectrum, are mitochondria-poor and primarily rely on glycolysis for energy production. Consequently, they are more prone to fatigue compared to oxidative muscles like the extensor digitorum longus and superficial quadriceps, which are better adapted for rapid, short-duration activities [21, 22]. Other fiber types, namely IIa and IIx, exhibit intermediate biophysical properties [19].

This study aims to screen a drug library using a PHF20/YY1 promoter reporter assay to identify potential drugs that can promote muscle differentiation. Additionally, this study investigates the in vivo efficacy of a selected drug candidate for its potential application in treating sarcopenia.

## Results

### Sulfasalazine Modulates YY1 Expression in Myoblasts

Our previous study demonstrated the necessity of YY1 for PHF20-mediated myogenic differentiation [14], with PHF20 augmenting YY1 transcription in C_2_C_12_ myoblasts. To identify drugs that enhance muscle differentiation through the PHF20-YY1 pathway, PHF20-pYY1-GFP and PHF20-pYY1-gLuc myoblast clones for high-throughput screening platform were established. The FDA-approved library was selected as it contains ∼1000 drugs that have already been approved in different medications in human, offering an appropriate library to identify drugs for repurposing. During the screening process, sulfasalazine emerged as a primary candidate that inhibits the YY1 promoter, as mediated by PHF20, in C_2_C_12_ cells (Fig. 1A). To ascertain the IC_50_ values of sulfasalazine on PHF20-mediated YY1 promoter inhibition, gaussia luciferase activity of PHF20-pYY1-gLuc C_2_C_12_ clones was measured after treatment in various concentrations of sulfasalazine. The calculated IC_50_ value for sulfasalazine was 24.3 μM (Fig. 1B). Furthermore, treatment with sulfasalazine decreased mRNA and protein levels of YY1 in normal C_2_C_12_ cells (Fig. 1C and 1D), providing evidence that sulfasalazine modulates YY1 expression in C_2_C_12_ cells.

**Figure 1.**
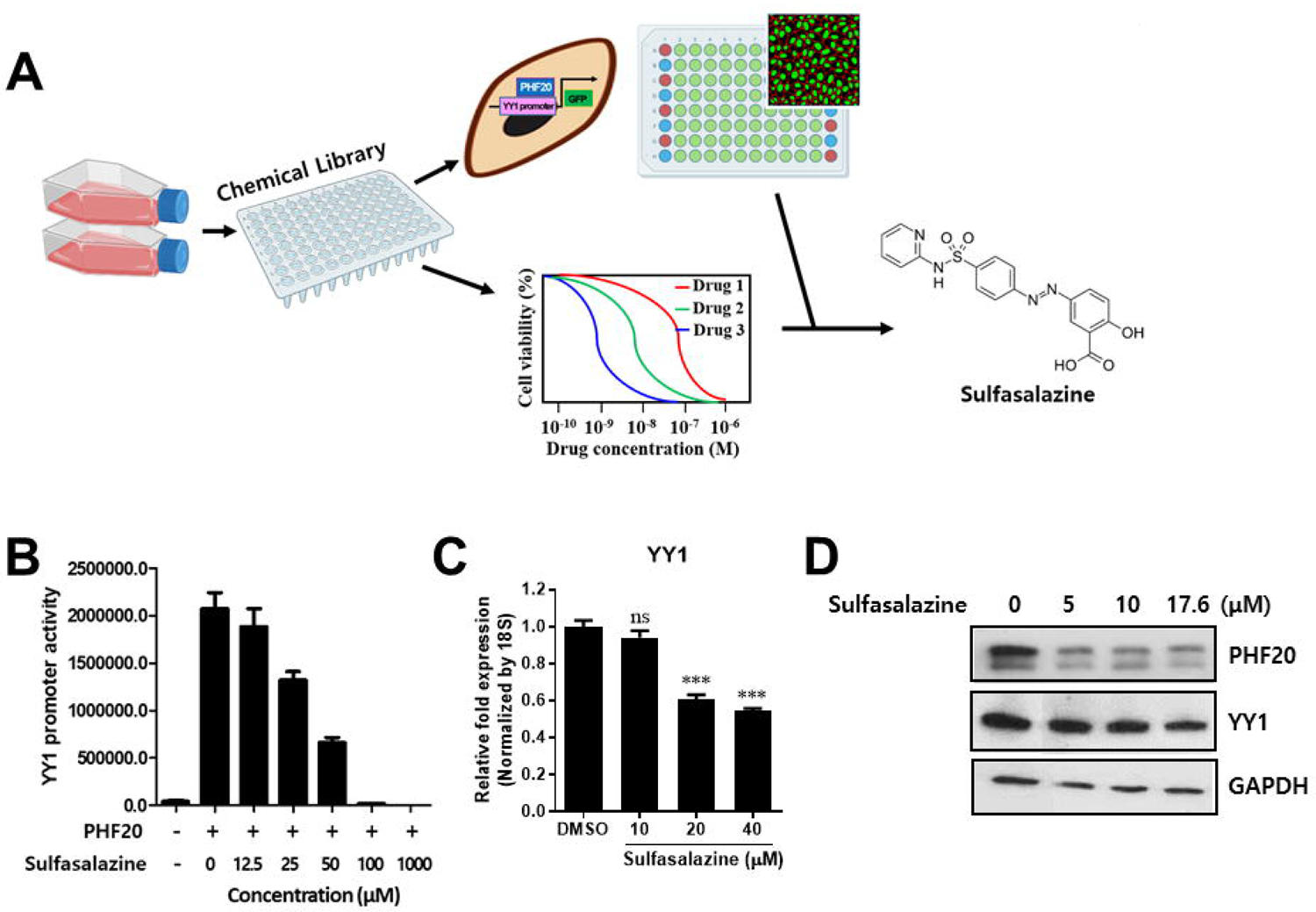
Cell-based screening of the approved drug library. (A) Schematic representation of the experimental process of PHF20-pYY1-GFP screening system. (B) YY1 promoter luciferase activity with PHF20 expression and sulfasalazine-dose gradient. Relative mRNA (C) and protein (D) expression of YY1 after dose-dependent treatment of sulfasalazine to C_2_C_12_ cells by qPCR and western blotting, respectively. Data represent the average of three replicates ± SEM. (***P < 0.001, ns indicates no significant difference, unpaired *t*-test).

### Sulfasalazine Enhances Myogenic Differentiation via Regulation of YY1 Expression

Prior research has indicated that YY1 may have an inhibitory effect on myogenesis by directly suppressing the expression of late-stage differentiation genes in skeletal muscle [11–13]. In order to further evaluate whether sulfasalazine, which downregulates YY1 expression, could encourage the myogenic differentiation of C_2_C_12_ cells, the gene expression of MYF5 and MYH as myogenic markers was measured by qPCR. MYF5 is typically expressed in proliferating myoblasts, and diminishes in differentiating myoblasts or myotubes [23]. MYH plays a critical role in muscle fiber formation and function during the late stages of myogenic differentiation. Intriguingly, sulfasalazine treatment led to decreased MYF5 expression, and increased MYH and myogenin expression during the myogenic differentiation process of C_2_C_12_ myoblasts (Fig. 2A and 2B). Additionally, MyHC expression in differentiated C_2_C_12_ cells with or without sulfasalazine was analyzed by using immunofluorescence staining. The morphometric analysis focused on myotube length, indicative of fiber size, as well as the fusion index and the number of myotubes, which are markers for the degree of myogenic differentiation. Cultures treated with sulfasalazine exhibited significant increases in fusion index, myotube length, the number of myotubes, and MyHC expression (Fig. 2C and 2D), demonstrating that sulfasalazine bolsters the myogenic differentiation of myoblasts (Fig. 2E).

**Figure 2.**
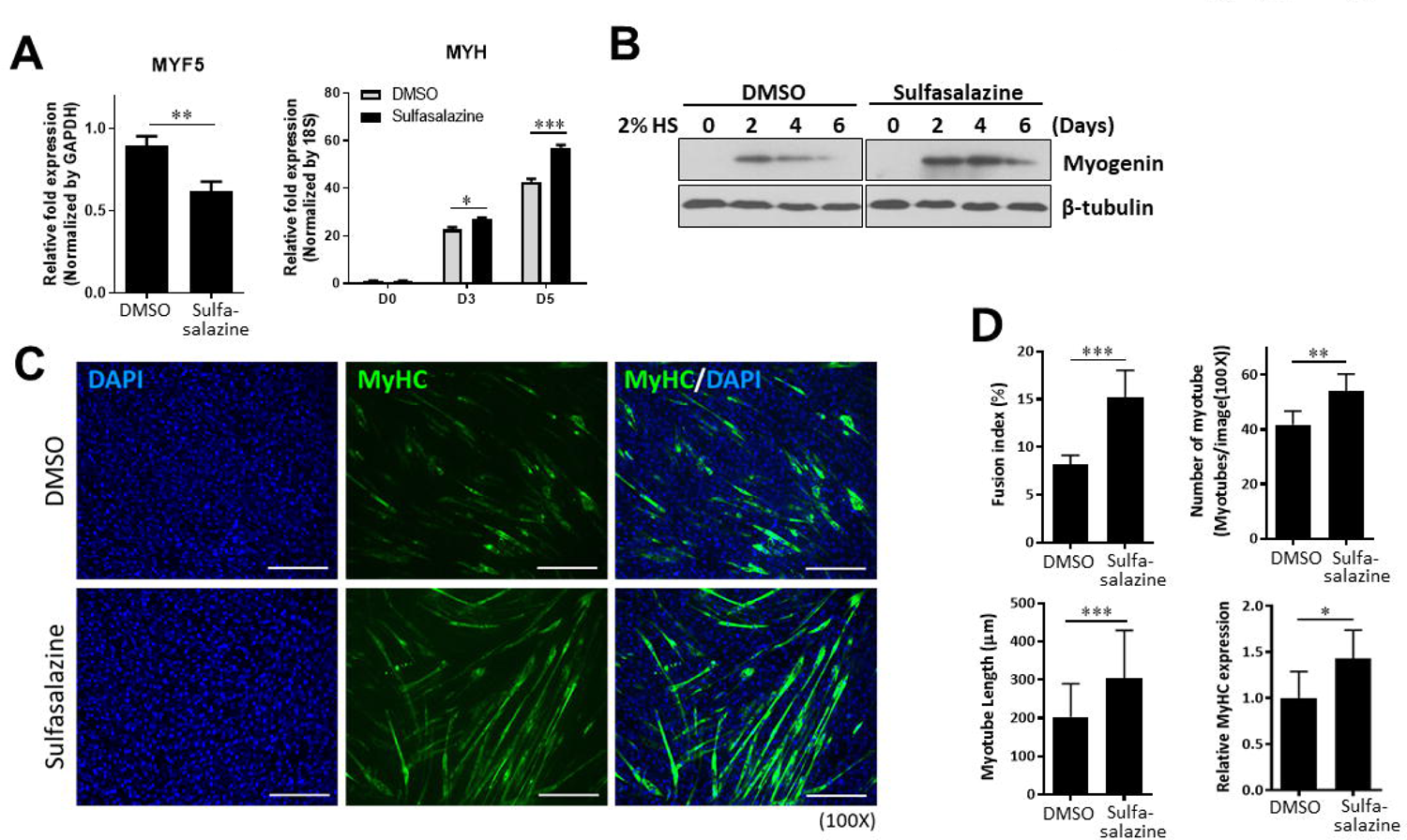
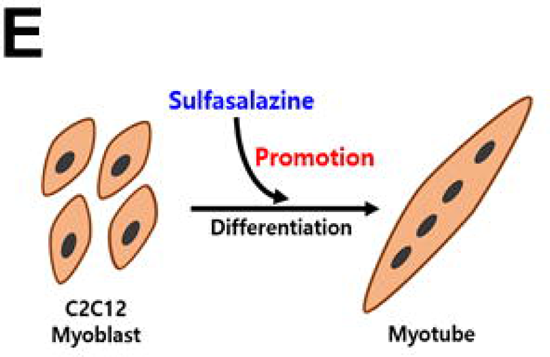
Sulfasalazine promotes myogenic differentiation through the regulation of YY1 expression. (A) Relative mRNA expression of *MYF5* and *MYH*, myogenic markers, in myogenic differentiated C_2_C_12_ cells with sulfasalazine-treatment by qPCR. Data represent the average of three replicates ± SEM. (B) Protein expression of myogenin in time-dependently myogenic differentiated C_2_C_12_ cells with sulfasalazine-treatment by western blotting. (C) The expression of MyHC (green) and nuclei (DAPI, blue) by immunofluorescence staining after myogenic differentiation of C_2_C_12_ cells with sulfasalazine treatment. Scale bar, 300 μm. (D) Morphometric analyses were performed on replicate samples (n=6). Fusion index, the number and length of myotubes, and MyHC expression were measured by Image J software. The fusion index was calculated as the ratio of the nuclei number in myocytes with two or more nuclei versus the total number of nuclei. (E) Illustration of the meaning of the experiment results. Data represent the average of 6 replicates ± SEM. D0, no differentiation; D3, differentiation for 3 days; D5, differentiation for 5 days; HS, horse serum. (***P < 0.001, **P < 0.01, *P < 0.05, unpaired *t*-test).

### Sulfasalazine Fosters a Shift from Oxidative Phosphorylation to Glycolysis in Myoblasts

Earlier studies have shown that myoblasts that have undergone myogenic differentiation exhibit a lower basal oxygen consumption rate (OCR), a measure of mitochondrial respiration, compared to proliferating myoblasts [24]. To investigate whether sulfasalazine-induced promotion of C_2_C_12_ cell differentiation involved a shift from oxidative phosphorylation to glycolysis in mitochondrial metabolism, OCR and extracellular acidification rate (ECAR) were monitored in C_2_C_12_ cells exposed to sulfasalazine. Consistent with previous reports, differentiated C_2_C_12_ cells under control conditions exhibited a lower basal OCR than undifferentiated cells (Fig. 3A). Sulfasalazine-treated C_2_C_12_ cells displayed a significantly lower OCR compared to control cells treated with DMSO (Fig. 3B and 3C), suggesting that sulfasalazine fosters a metabolic shift from oxidative phosphorylation to glycolysis (Fig. 3D). Moreover, sulfasalazine increased the basal ECAR (Fig. 3E). PGC-1α is renowned as a central regulator of mitochondrial biogenesis [25] and orchestrates mitochondrial biogenesis and respiration, ensuring metabolic homeostasis [26, 27]. To further assess the involvement of PGC-1α in the sulfasalazine-mediated reduction in OCR, the changes of gene expression of *PGC1α* was measured in sulfasalazine-treated C_2_C_12_ cells. It was confirmed that sulfasalazine treatment led to a dose-dependent decrease in PGC-1α expression (Fig. 3F), suggesting that the metabolic shift from oxidative phosphorylation to glycolysis, induced by sulfasalazine, can be attributed to a reduction in PGC-1α expression.

**Figure 3.**
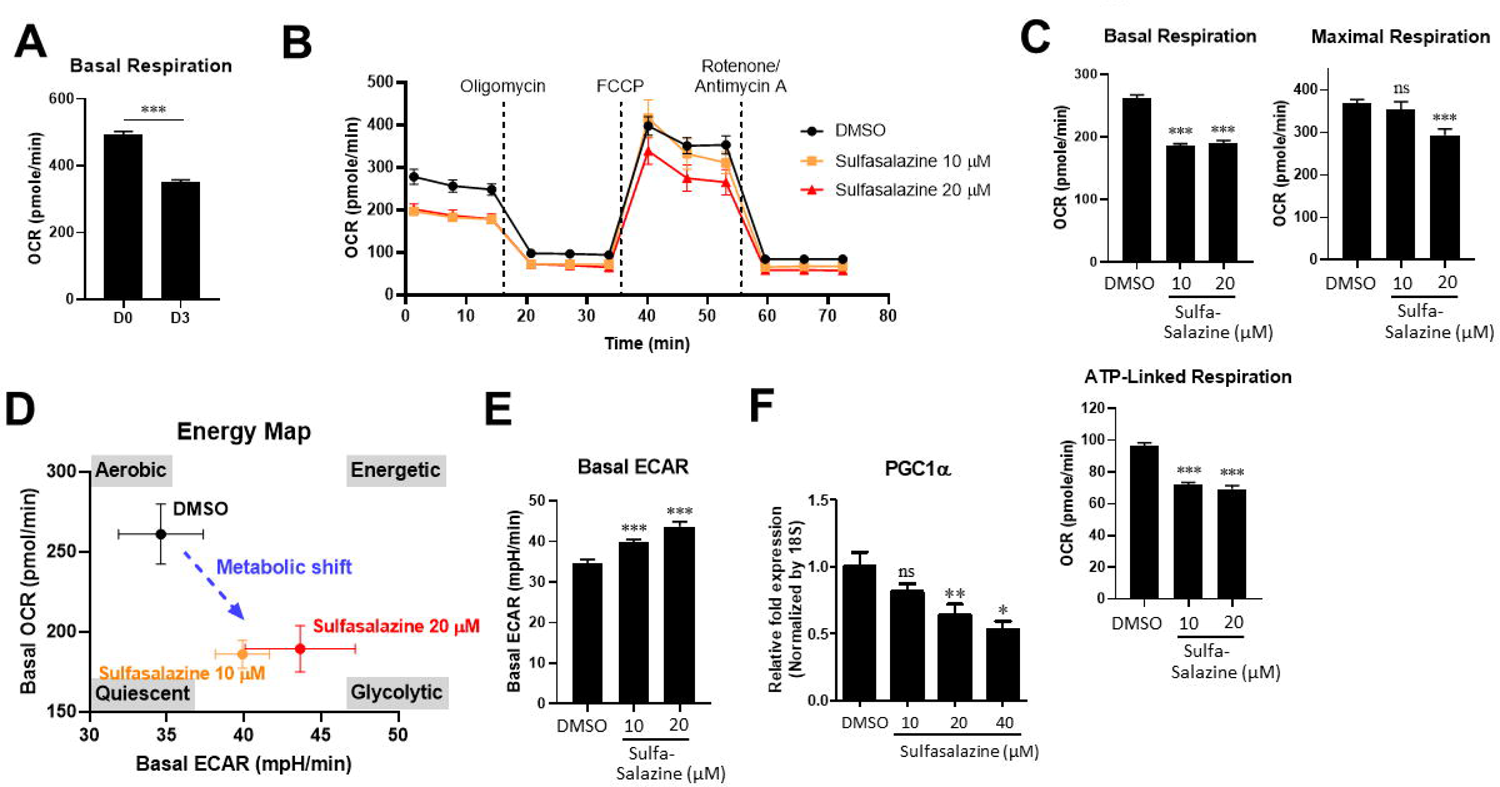
Sulfasalazine increases glycolysis in C_2_C_12_ myoblasts. (A) Comparison of basal OCR between myoblast and myogenic differentiation for 3 days of C_2_C_12_ cells. (B) OCR measurements were obtained over time (min) using an extracellular flux analyzer (Seahorse Bioscience). Mitochondrial respiration reflected by OCR levels was detected in C_2_C_12_ cells with sulfasalazine treatment under basal conditions or following the addition of oligomycin (1.5 μM), the uncoupler FCCP (0.5 μM), and a combination of rotenone (0.5 μM) and antimycin A (0.5 μM) (n = 5). (C) The rates of basal respiration, ATP-linked respiration, and maximal respiratory capacity were quantified by normalization of OCR level to the total protein OD values. (D) Energy map for cells cultured under the two-dose conditions of sulfasalazine. Data are expressed as mean ± SEM (n = 5). (E) Basal ECAR under basal conditions in C_2_C_12_ cells with sulfasalazine. Data represent the average of four replicates ± SEM. Data represent the average of four replicates ± SEM. All of Seahorse raw data were analyzed using Seahorse Wave Software (version 2.3.0.19). (F) qPCR analysis of *PGC1α* mRNA expression in response to sulfasalazine treatment. Data represent the average of three replicates ± SEM. (***P < 0.001, ns indicates no significant difference, unpaired *t*-test).

### Sulfasalazine Promotes the Expression of Fast Myosin Heavy Chain through a Metabolic Shift to Glycolysis

In skeletal muscles, the composition of muscle fiber types dictates their varied properties, such as contractility (slow-twitch or fast-twitch) and fatigue resistance, as well as metabolic characteristics (oxidative or glycolytic) [28]. Additionally, it has been documented that PGC1α plays a role in muscle fiber-type switching, favoring an increased proportion of oxidative fiber types [29]. To investigate whether sulfasalazine influences fiber-type composition due to the decrease in PGC1α, the gene expression for MyHC isoforms was monitored after myogenic differentiation of C_2_C_12_ cells with or without sulfasalazine treatment. Sulfasalazine treatment was found to enhance the mRNA and protein expression of genes associated with the glycolytic fast-twitch muscle marker (fast MyHC) in myogenic differentiated C_2_C_12_ cells (Fig. 4A and 4B). Notably, there was a significant increase in the expression of MyHC type II (MYH2 for IIa, MYH1 for IIx, and MYH4 for IIb), which are glycolytic fast fiber types, in myogenic differentiated C_2_C_12_ cells treated with sulfasalazine compared to DMSO-treated control cells. However, expression of the oxidative slow fiber type (MYH7 for MyHC type I) did not show a significant increase (Fig. 4A). Additionally, an increased expression of MyHC type IIb, a representative glycolytic fiber type, was observed in myotube-differentiated C_2_C_12_ cells treated with sulfasalazine (Fig. 4B). These findings support the hypothesis that sulfasalazine modulates fiber-type pathways (Fig. 4C).

**Figure 4.**
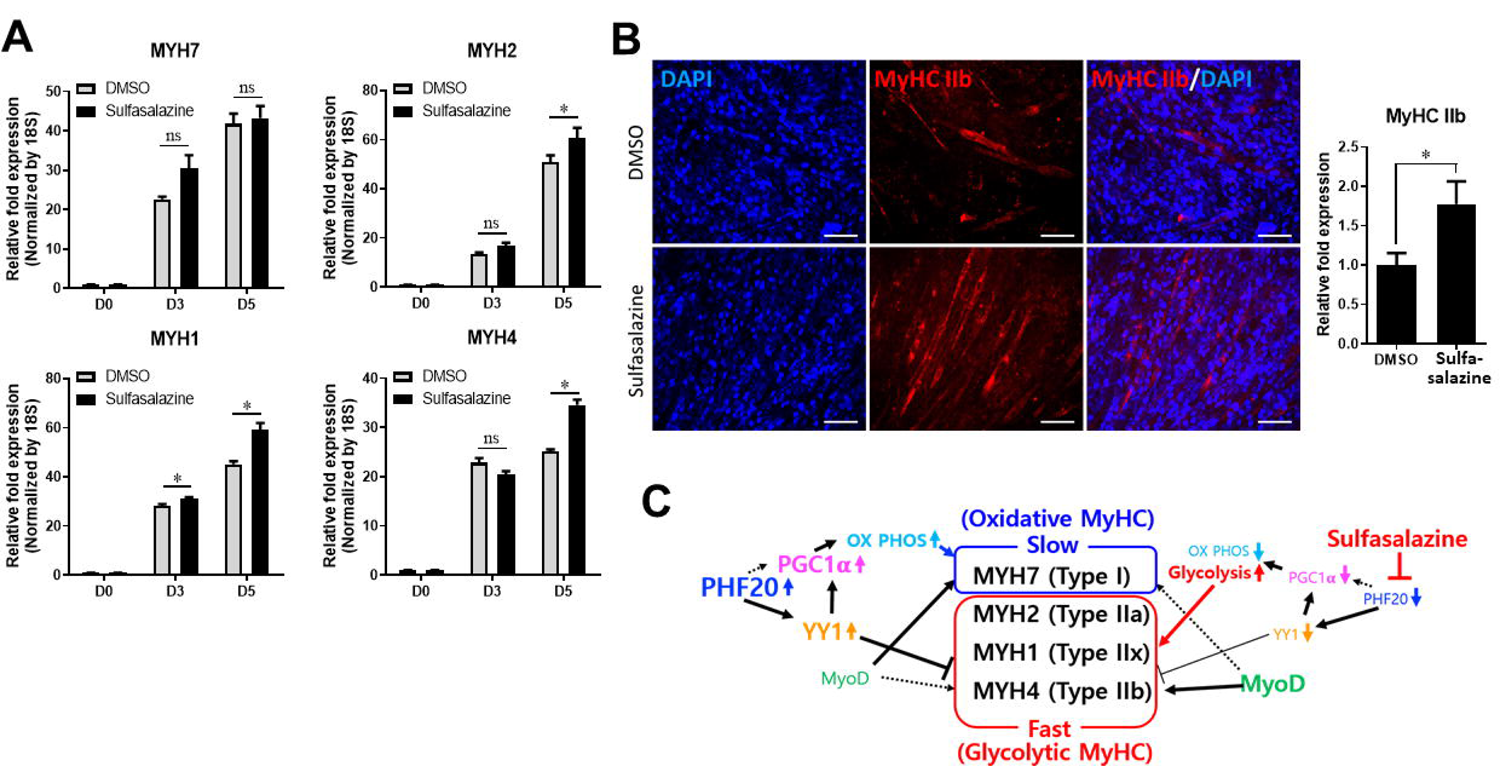
Sulfasalazine promotes the expression of fast type-myosin heavy chains. (A) MyHC type mRNA expression at day 3 and day 5 of C_2_C_12_ cell differentiation with sulfasalazine. mRNA of *MYH7* for type I, *MYH2* for IIa, *MYH1* for IIx, and *MYH4* for IIb were analyzed by qPCR. Data represent the average of three replicates ± SEM. (B) Immunofluorescence staining of the MyHC type IIb (red) and the nuclei (DAPI, blue) were compared between the myogenic differentiated C_2_C_12_ cells treated with DMSO and sulfasalazine. Quantification of this expression is indicated on the right panel. The expression level of MyHC type IIb was quantified using Image J software. Scale bar, 100 μm. (C) Schematic of signals related to the promotion of the myogenic differentiation by sulfasalazine. D0, no differentiation; D3, differentiation for 3 days; D5, differentiation for 5 days. (*P < 0.05, ns indicates no significant difference, unpaired *t*-test).

### Sulfasalazine Enhances Physical Performance in Sarcopenic Animal Models through Fiber Type Switching to Glycolytic Fast Fiber

Skeletal muscle, as the largest and most regenerative organ in mammals, can experience functional decline due to aging. The age-associated loss of skeletal muscle mass can lead to physical dysfunction and imbalance, particularly increasing the risk of sarcopenia [30, 31]. To explore whether sulfasalazine can enhance physical performance in vivo by promoting muscle differentiation and the expression of specific muscle fiber types, sulfasalazine treatment was performed in animal models. Various animal models were prepared to emulate the human sarcopenia or muscle loss. Aged mice were used to model geriatric diseases (Fig. 5), along with hindlimb disuse (Fig. 6) and hindlimb damage models (Fig. 7) to simulate human leg muscle injury. To model human sarcopenic obesity, a high-fat diet (HFD)-fed mouse model with hindlimb damage was employed (Fig. 8). Sarcopenia and muscle loss primarily occur in the elderly; thus, 50-week-old male mice were used. To simulate human sarcopenia muscle loss, muscle damage was induced in one hindlimb of the mice using cardiotoxin injection or Velcro fixation.

**Figure 5.**
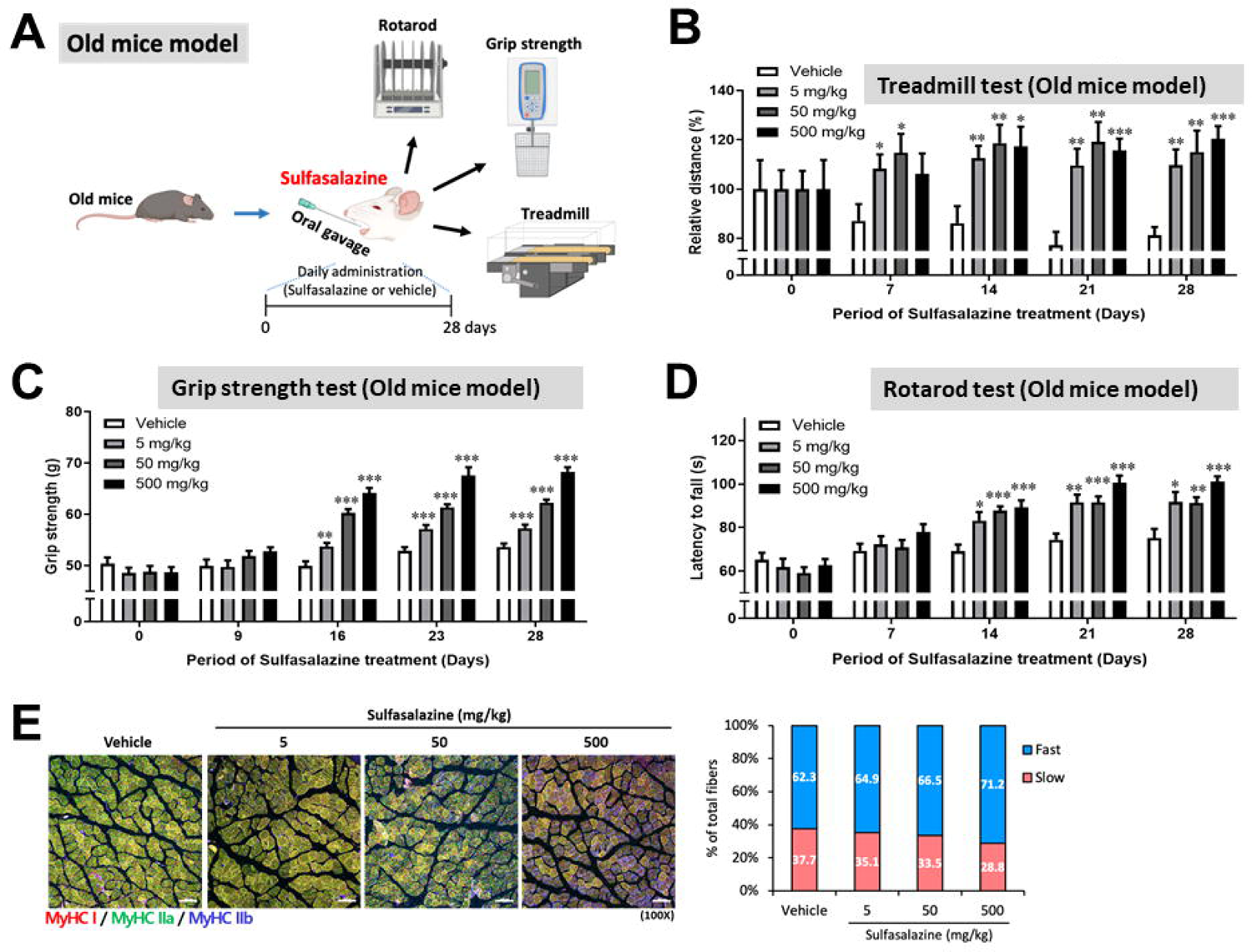
Sulfasalazine improves physical performance of old mice model. 50-week-old male mice were used for this experiments as aged mice. (A) Schematic representation of the experimental process of the effects of sulfasalazine on physical performance in old mice model. (B) The running endurance of the mice was evaluated by a treadmill test. (C) The grip strength of the hindlimbs was measured at regular intervals during administration. (D) Vehicle- and sulfaslalzine-treated mice performed equally well during the first three trials of the accelerated rotarod motor test. Statistical significance was determined using *t*-test between the vehicle-control group and each sulfasalazine administration group. (E) Immunofluorescence staining of MyHC type I (red), IIa (green), and IIb (blue). Quantification of each MyHC type is indicated on the right panel. Scale bar, 100 μm. Treadmill data represent the average of same groups ± SEM. Rotarod data represent the average of values repeated 3 times per mouse in each group ± SEM. Grip data represent the average of values repeated 5 times per mouse in each group ± SEM. (n = 6, ***P < 0.001, **P < 0.01, *P < 0.05, unpaired *t*-test).

**Figure 6.**
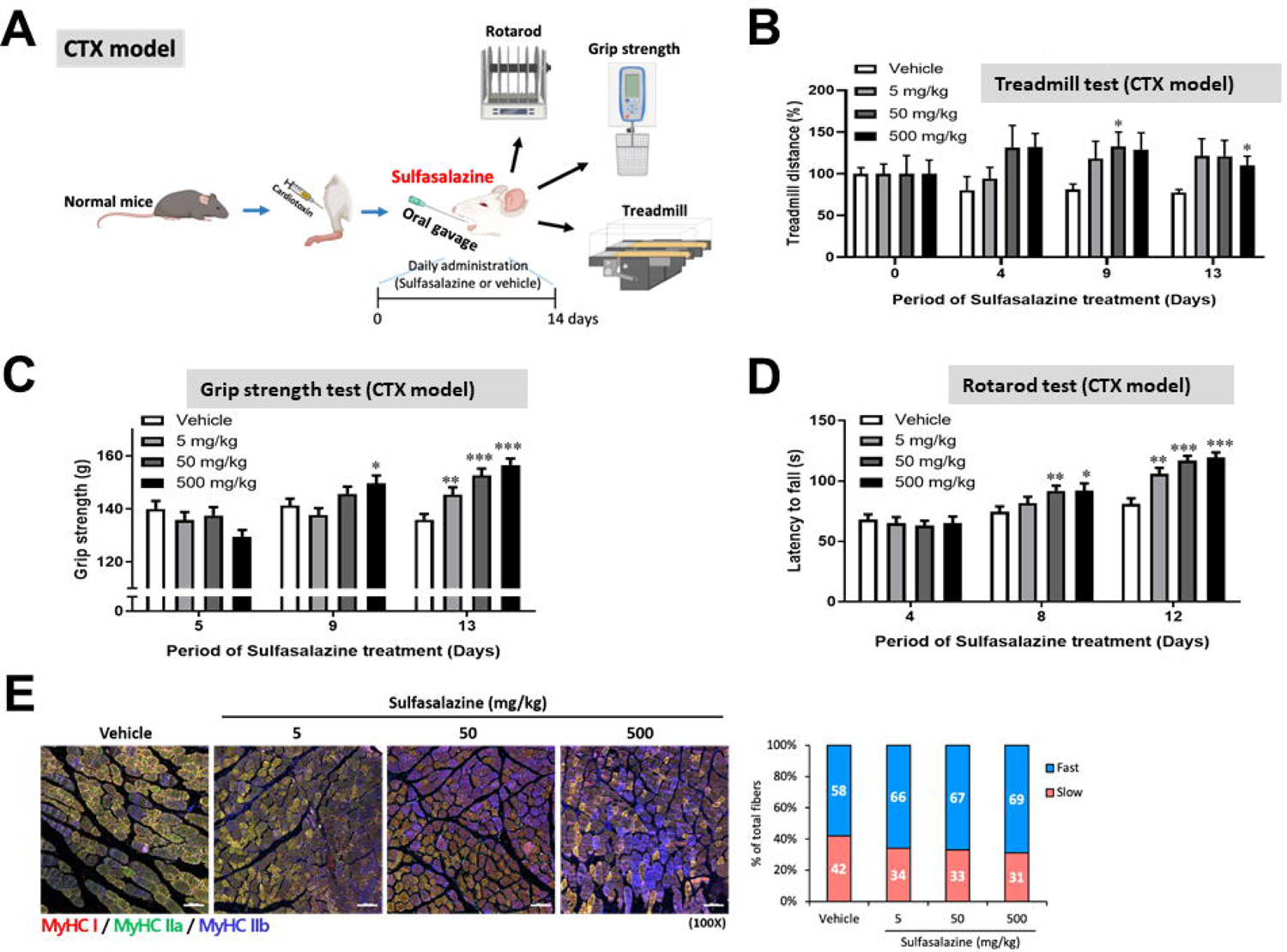
Sulfasalazine improves physical performance of CTX model. Muscle damage was induced in one of hindlimb of mice using cardiotoxin to embody muscle loss in human sarcopenia. (A) Schematic representation of the experimental process of the effects of sulfasalazine on physical performance in CTX model. The treadmill running test (B), the grip strength test of the hindlimbs (C), and rotarod test (D) of vehicle- and sulfaslalzine-treated mice performed appropriately. Statistical significance was determined using *t*-test between the vehicle-control group and each sulfasalazine administration group. (E) Immunofluorescence staining of MyHC type I (red), IIa (green), and IIb (blue). Quantification of each MyHC type is indicated on the right panel. Scale bar, 100 μm. Treadmill data represent the average of same groups ± SEM. Rotarod data represent the average of values repeated 3 times per mouse in each group ± SEM. Grip data represent the average of values repeated 5 times per mouse in each group ± SEM. (n = 6, ***P < 0.001, **P < 0.01, *P < 0.05, unpaired *t*-test).

**Figure 7.**
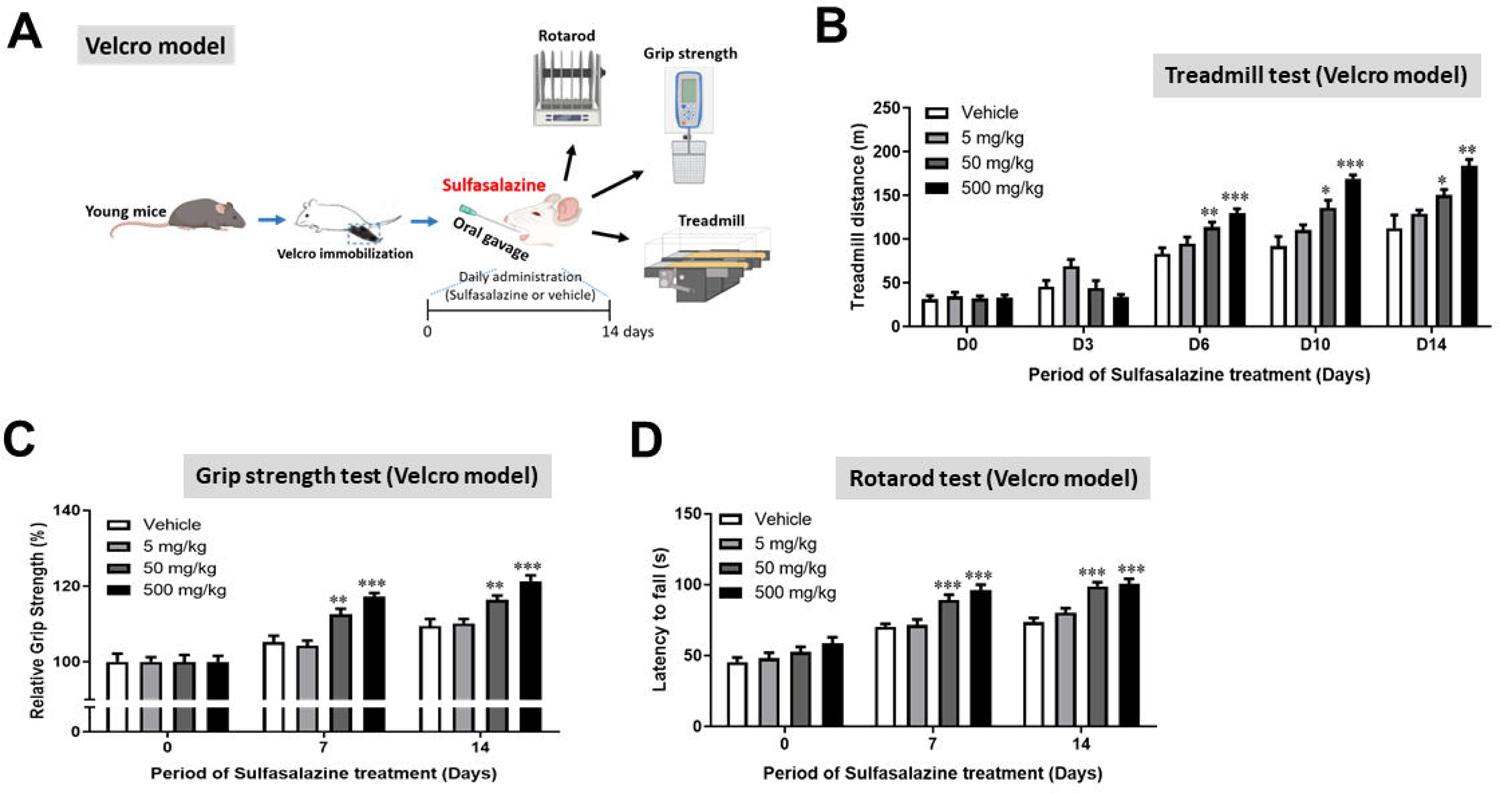
Sulfasalazine improves physical performance of Velcro model. Muscle damage was induced in one of hindlimb of mice using Velcro immobilization to embody muscle loss in human sarcopenia. (A) Schematic representation of the experimental process of the effects of sulfasalazine on physical performance in Velcro model. The treadmill running test (B), the grip strength test of the hindlimbs (C), and rotarod test (D) of vehicle- and sulfaslalzine-treated mice performed appropriately. Statistical significance was determined using *t*-test between the vehicle-control group and each sulfasalazine administration group. Treadmill data represent the average of same groups ± SEM. Rotarod data represent the average of values repeated 3 times per mouse in each group ± SEM. Grip data represent the average of values repeated 5 times per mouse in each group ± SEM. (n = 6, ***P < 0.001, **P < 0.01, *P < 0.05, unpaired *t*-test).

**Figure 8.**
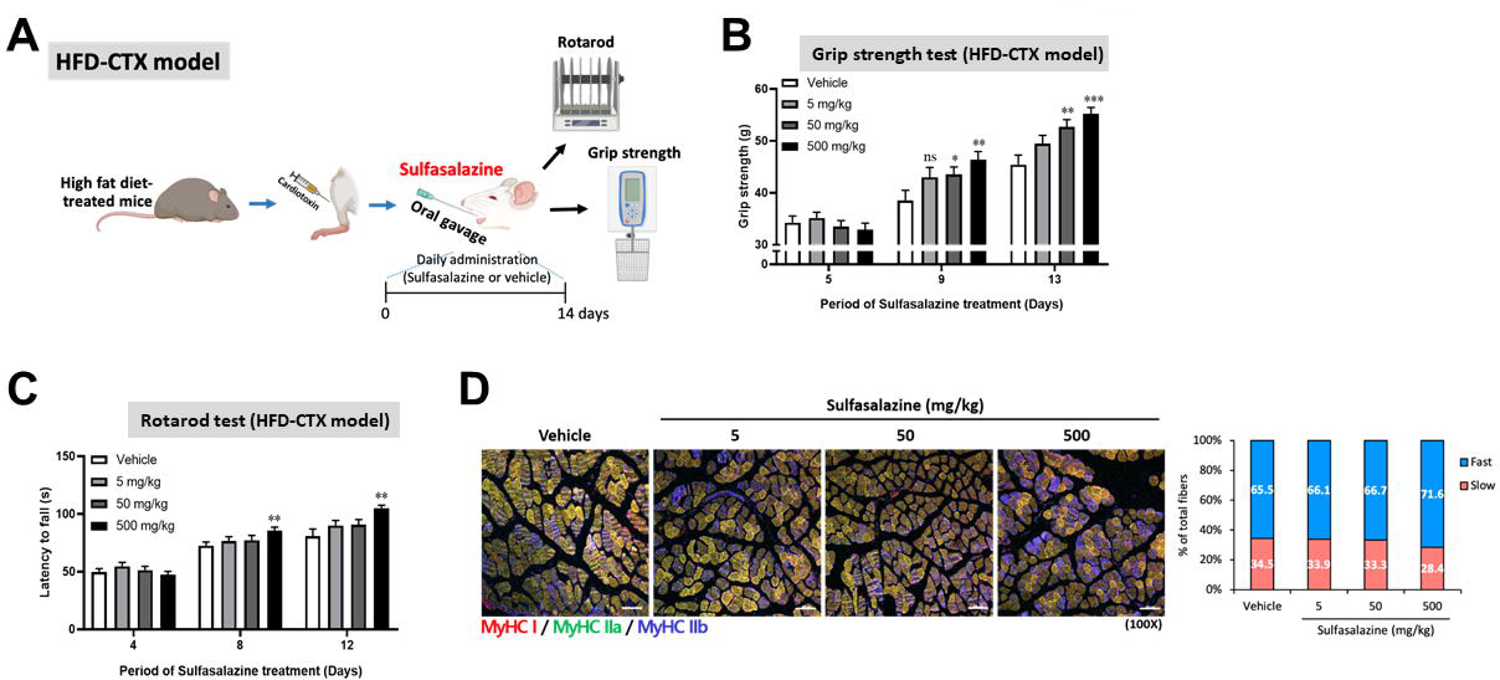
Sulfasalazine improves physical performance of HFD-CTX model. Muscle damage was also induced in one of hindlimb of obese mice fed a HFD for 19 weeks using cardiotoxin to embody muscle loss in human sarcopenic obesity. (A) Schematic representation of the experimental process of the effects of sulfasalazine on physical performance in HFD-CTX model. The grip strength test of the hindlimbs (B), and rotarod test (C) of vehicle- and sulfaslalzine-treated mice performed appropriately. Statistical significance was determined using *t*-test between the vehicle-control group and each sulfasalazine administration group. (D) Immunofluorescence staining of MyHC type I (red), IIa (green), and IIb (blue). Quantification of each MyHC type is indicated on the right panel. Scale bar, 100 μm. Rotarod data represent the average of values repeated 3 times per mouse in each group ± SEM. Grip data represent the average of values repeated 5 times per mouse in each group ± SEM. (n = 6, ***P < 0.001, **P < 0.01, *P < 0.05, ns indicates no significant difference, unpaired *t*-test).

Physical performance tests such as treadmill, rotarod, and grip strength tests were conducted weekly on a staggered schedule. Sulfasalazine was orally administered daily to aged mice (Old mice model) for 28 days, hindlimb-damaged mice by cardiotoxin (CTX model) for 14 days, hindlimb-immobilized mice by Velcro (Velcro model) for 14 days, and high-fat diet-fed mice with hindlimb damage (HFD-CTX model) for 14 days. Physical performance tests were conducted at suitable intervals during the sulfasalazine administration period. The treadmill test, rotarod test, and grip strength test were used to evaluate the effect of sulfasalazine on physical performance in each animal model, with the overall experimental design illustrated in the respective figures (Fig. 5A, 6A, 7A, and 8A). Sulfasalazine-treated groups demonstrated significantly longer treadmill distances than the vehicle-control group throughout the experiment (Fig. 5B, 6B, and 7B). Additionally, in the hindlimb grip tests, sulfasalazine-treated groups exhibited markedly improved maximal muscle strength compared to vehicle-controls in a dose- and time-dependent manner (Fig. 5C, 6C, 7C, and 8B). In the accelerating rotarod test (acceleration from 4 to 40 rpm in 5 min), mice treated with sulfasalazine showed notably enhanced performance compared to control animals (Fig. 5D, 6D, 7D, and 8C). To understand the underlying mechanism for the improvement in exercise performance, the MyHC type of the muscle fiber of all mice was analyzed by immunofluorescence. As depicted in Figures 5E, 6E, and 8D, fast fiber types (specifically MyHC type IIb) were increased in a dose-dependent manner in the sulfasalazine-treated groups, consistent with the in vitro findings in Figure 4. The switching of fiber type from slow to fast in muscle tissue is a likely contributor to the enhanced physical performance of sulfasalazine-treated groups compared to the control group. In summary, sulfasalazine treatment improved physical performance in various muscle loss models by inducing a switch from slow to fast fiber types in muscles.

### Damaged Muscle Fibers are Restored Through the Increase in Satellite Cells by Sulfasalazine Treatment

To delve further into the impact of sulfasalazine on physical performance, alterations in muscle fibers were monitored in mice. As anticipated, muscle fibers suffered damage due to Velcro immobilization (Fig. 9A). During the spontaneous healing of muscle fibers impaired by Velcro immobilization, sulfasalazine administration notably enhanced the recovery process (Fig. 9B and 9C). Remarkably, a considerable number of cells expressing PAX7, a satellite cell marker, were identified in muscle tissue treated with sulfasalazine (Fig. 9B and 9C). These findings strongly support the hypothesis that sulfasalazine administration facilitates myogenic differentiation and increases the number of satellite cells expressing PAX7, which in turn leads to improved physical performance in the treated mice.

**Figure 9.**
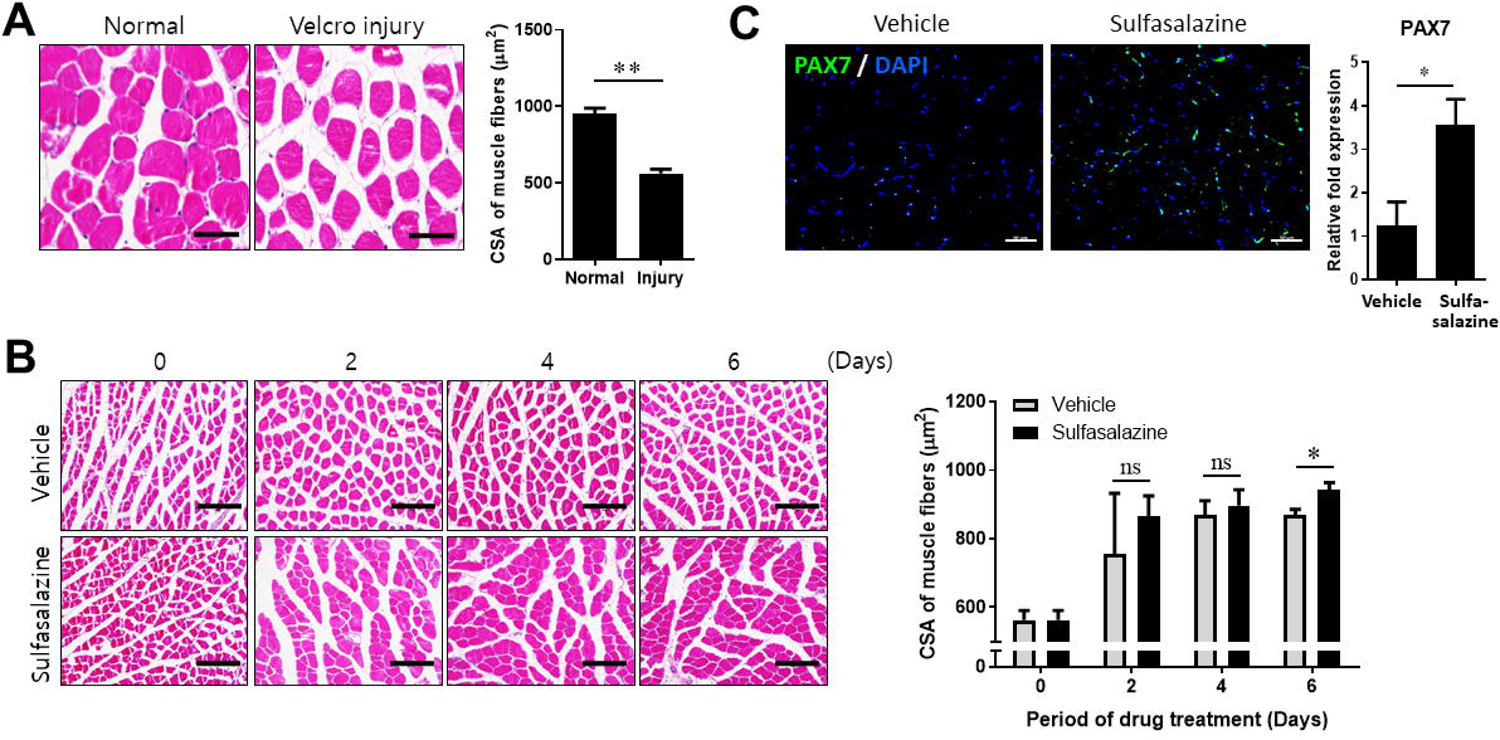
Damaged muscle fibers are restored by sulfasalazine treatment. Muscle damage was induced in one of hindlimb of mice using Velcro immobilization. (A) Comparison of H&E staining of normal and damaged GA muscle tissues. Average of muscle fiber cross-sectional area (CSA) was measured by Image J software. Quantification of each size of muscle fibers is indicated on the right panel. Scale bar, 50 μm. (B) H&E staining for the effects of promoting recovery of damaged muscle tissues by sulfasalazine. Quantification of each size (CSA) of muscle fibers is indicated on the right panel. Scale bar, 150 μm. (C) Immunofluorescence staining of PAX7 (green), a marker of muscle satellite cells, and the nuclei (DAPI, blue). Quantification of PAX7 expression is indicated on the right panel. Scale bar, 50 μm. (**P < 0.01, *P < 0.05, ns indicates no significant difference, unpaired *t*-test).

## Discussions

Patients with sarcopenia experience a slow gait and reduced muscular endurance, which makes daily life challenging and often necessitates help from others. Additionally, they are prone to osteoporosis, falls, and fractures [32]. With the decrease in muscle mass, the blood and hormone buffering capacity of muscles diminishes, leading to a lower basal metabolic rate. Consequently, managing chronic diseases becomes difficult, with diabetes and cardiovascular diseases tending to worsen. As a result, sarcopenia has recently been recognized as a geriatric disease worldwide. Sarcopenia is characterized by the loss of muscle mass associated with aging [3]. Though muscle mass reduction due to aging can be viewed as a natural process, the age at which sarcopenia is diagnosed has been decreasing. Furthermore, no new drugs have yet been approved by the FDA for the treatment of sarcopenia [33]. While companies worldwide initiated research into sarcopenia treatments before it was officially recognized as a disease, currently, no company is pursuing the development of a new drug. Developing a therapeutic agent proves challenging as muscle loss with aging does not stem from a single biological mechanism. Moreover, an effective therapeutic agent must address not only muscle mass but also physical performance [34].

Sulfasalazine, an anti-inflammatory drug with over 50 years of usage, is primarily prescribed for conditions such as early-stage rheumatism and inflammatory bowel diseases, including Crohn’s disease and ulcerative colitis. Although the precise mechanism of sulfasalazine remains unclear, it is known to inhibit the transcription factor nuclear factor kappa-B (NF-κB), suppressing the transcription of NF-κB responsive pro-inflammatory genes, including TNF-α [35]. Additionally, sulfasalazine inhibits TNF-α expression by inducing caspase-8-mediated apoptosis in macrophages [36], and prevents osteoclast formation by suppressing the expression of receptor activator of NF-κB ligand (RANKL) and stimulating osteoprotegerin, a natural inhibitor of RANKL [37]. This study introduced a novel mechanism where sulfasalazine regulates PHF20-mediated YY1 expression (Fig. 1). Specifically, the study emphasized the transcriptional and translational regulation of YY1 due to sulfasalazine treatment. The observed reduction in YY1 expression upon sulfasalazine treatment is speculated to hinder formation of the YY1/Ezh2/HDAC complex, which suppresses expression of MCK and MyHC DNA, ultimately promoting expression of late-stage differentiation genes by serum response factor and MyoD. YY1 is known to regulate PGC1α, a critical regulator of mitochondrial biogenesis [25–27]. The study confirmed that sulfasalazine-induced reduction in YY1 expression leads to decreased PGC1α expression, thereby promoting a metabolic shift towards glycolysis in the mitochondrial metabolism of C_2_C_12_ myoblasts.

It is well-documented that a reduction in muscle mitochondria triggers a slow-to-fast fiber transition and a shift to a glycolytic energy production system [17]. This study further established that sulfasalazine induces a metabolic shift during the myogenic differentiation of C_2_C_12_ myoblasts, increasing gene expression of glycolytic fast fiber type (Fig. 3). Fast type MyHC was also observed to increase in muscle tissue following administration of sulfasalazine in various sarcopenic animal models (Fig. 5, 6, and 8). Notably, in the Velcro model, one of the animal models for sarcopenia, sulfasalazine administration was found to promote the recovery of muscle tissue damaged by Velcro immobilization (Fig. 9B). Interestingly, an increase in the population of satellite cells, which are muscle stem cells, was observed post sulfasalazine administration (Fig. 9C). This suggests that sulfasalazine enhances muscle regeneration in the Velcro model through the elevation of satellite cells, and promotes myogenic differentiation of these cells, facilitating muscle tissue recovery. This study demonstrates that sulfasalazine fosters myogenic differentiation by augmenting glycolytic fast muscle fibers through the mitochondrial metabolic reprogramming of C_2_C_12_ myoblasts. This, in turn, enhances the physical performance of animal models through the increase of fast muscle fibers and stimulates the proliferation of satellite cells, contributing to the recovery of damaged muscle tissue.

The study employed a daily dose of 50 mg/kg of sulfasalazine in mice. Extrapolating to human dosage, it was determined that a 60 kg adult could be administered 240 mg once a day [38]. This dosage is lower than the recommended dosage for ulcerative colitis (500∼1000 mg every 6 to 8 hours per day) or rheumatoid arthritis (500∼1000 mg once or twice a day in divided doses) in adults. Thus, a daily dose of 240 mg of sulfasalazine is considered safe and potentially suitable for treating human sarcopenia. The exact mechanisms through which sulfasalazine directly regulates PHF20-mediated YY1 and promotes proliferation of satellite cells remain to be elucidated. Nonetheless, based on the findings of this study, sulfasalazine emerges as a promising candidate for the treatment of sarcopenia.

## Materials and Methods

### Cell culture

C_2_C_12_ myoblast cells (ATCC, CRL-1772) were maintained in DMEM (WELGENE, Korea) supplemented with 10% fetal bovine serum (FBS) (WELGENE) and 1% Anti-anti (Gibco) at 37 °C in 5% CO2. Differentiation was initiated 24 h after seeding by changing to differentiation medium consisting of DMEM supplemented with 2% horse serum (HS) and 1% Anti-anti. The medium was changed for every 2 days.

### Establishment of C_2_C_12_/Tet-On-inducible PHF20/pYY1 reporter cells

C_2_C_12_ cells were treated with lentivirus-TET3G and selected with G418 (1.2 mg/ml) to generate C_2_C_12_-TET3G myoblast cells. Next, C_2_C_12_-TET3G cells were transiently transfected with pLVX-TRE3G-PHF20 and selected with puromycin (3 μg/ml) for Tet-On inducible PHF20 cells. The expression of PHF20 was confirmed in the media with doxycycline (Doxy). The antibiotic resistance gene against puromycin in a GFP or gaussia luciferase reporter plasmid (pEZX-LvPF02 or pEZX-LvPG02, GeneCopoeia) containing a YY1 promoter sequence was changed to hygromycin resistance gene. And then each of these plasmids was introduced into C_2_C_12_/Tet-On-inducible PHF20 cells. Stable cells were selected using hygromycin (250 μg/mL) and kept frozen before use.

### Drug screening

The stable C_2_C_12_/Tet-On-inducible PHF20/YY1 reporter-GFP cells were treated with drugs from an FDA-approved drug library (Selleckchem.com) at 10 μM for 24 h right after treating with doxy for 48 h. GFP intensity of cells treated with each drug library was measured by the fluorescence microreader (GloMax Discover, Promega). The inhibition rate for GFP intensity was calculated, and hit was selected by sequentially arranging them according to the degree of inhibition.

### Immunoblot analysis

Western blotting was performed as previously described [39, 40]. After the comp0letion of experimental conditions, cells were placed on ice and extracted with lysis buffer, cells were placed on ice and extracted with lysis buffer containing 50 mM Tris-HCl, pH7.5, 1% v/v Nonidet P-40, 120 mM NaCl, 25 mM sodium fluoride, 40 mM β-glycerol phosphate, 0.1 mM sodium orthovanadate, 1 mM phenylmethylsulfonyl fluoride, 1 mM benzamidine, and 2 μM microcystin-LR. Lysates were centrifuged for 30 min at 13,000 rpm. The cell extracts were resolved by 10.0–7.5% SDS-PAGE, and transferred to Immobilon-P membranes (Millipore). The filters were blocked for 1 h in 1× tri-buffered saline buffer (140 mM NaCl, 2.7 mM KCl, and 250 mM Tris-HCl, pH7.4), containing 5% skim milk and 0.2% Tween-20, followed by an overnight incubation with each primary antibody diluted 1000 folds at 4 °C. Antibodies for myogenin (DSHB, F5D), YY1 (Santa Cruz, sc-7341), and PHF20 (Cell Signaling Technology, #3934) were used as a primary antibody. The secondary antibody was horseradish peroxidase-conjugated anti-mouse IgG or anti-rabbit IgG (Koma biotech, Seoul, Korea), diluted 2000-fold in the blocking buffer. The detection of protein expression was visualized by enhanced chemiluminescence, according to the manufacturer’s instructions.

### Real-time quantitative reverse transcription polymerase chain reaction (qRT-PCR)

Total RNA was extracted using TRIzol reagent (Invitrogen). cDNA was synthesized using the manufacturer’s instructions for SuperScript II (Invitrogen, 18064022). The reactions were up in triplicate with the GoTaq qPCR Master Mix (Promega) and run on AriaMx Real-time PCR (Agilent). The sequences of the primers that were used for the qPCR were as follows; mouse GAPDH, forward (5′-TCCAGTATGACTCCACTC-3′), reverse (5′-ATTTCTCGTGGTTCACAC-3′); mouse YY1, forward (5′-CAGAAGCAGGTGCAGATC AGACCCT-3′), and reverse (5′-GCACCACCACCCACGGAATCG-3′); mouse MYF5, forward (5′-AAGGCTCCTGTATCCCCTCAC-3′) and reverse (5′-TGACCTTCTTCAGGCGTCTAC-3′); mouse MYH, forward (5′-ACTGTCAACACTAAGAGGGTCA-3′) and reverse (5′-TTGGATGATTTGATCTTCCAGGG-3′); mouse PGC1α, forward (5′-CGGAAATCATATCCAACCAG-3′) and reverse (5′-TGAGGACCGCTAGCAAGTTTG-3′); mouse MYH7, forward (5′-ATGAGCTGGAGGCTGAGCA-3′), and reverse (5′-TGCAGCCGCAGTAGGTTCTT-3′); mouse MYH2, forward (5′-ATTCTCAGGCTTCAGGATTTGGTG-3′), and reverse (5′-CTTGCGGAACTTGGATAGATTTGTG-3′); mouse MYH1, forward (5′-AAGGGTCTGCGCAAACATGA-3′), and reverse (5′-TTGGCCAGGTTGACATTGGA-3′); mouse MYH4, forward (5′-GAGTTCATTGACTTCGGGATGG-3′), and reverse (5′-TGCTGCTCATACAGCTTGTTCTTG-3′).

### Immunofluorescence in cells

C_2_C_12_ cells was myogenic differentiated for 5 days using differentiation medium (DMEM including 2% HS). Immunofluorescence staining was performed by fixing C_2_C_12_ myotubes in 4% paraformaldehyde for 20 min, permeabilized with 0.2% Triton X-100 for 15 min and blocked with 2% bovine serum albumin in PBS for 30 min. Myotubes were incubated with antibodies against MyHC (DSHB, MF20) or MyHC type IIb (DSHB, BF-F3) overnight at 4 °C, followed by antibodies conjugated with Alexa Fluor 488 (Invitrogen, A-11001) for 1 h. The nuclei were counterstained with DAPI. Morphometric analyses were performed using Image J software.

### Measurement of OCR

An Agilent Seahorse XF Cell Mito Stress Kit was used to measure OCR. Briefly, C_2_C_12_ myoblasts (1.2×10^4^/well) were plated on the Seahorse 96-well culture plate. After allowing the cells to acclimate in a 37 °C non-CO2 incubator, the analyzer was calibrated, and the OCR was measured through sequential injection of electron transport chain inhibitors: 1.5 μM oligomycin (ATP synthase inhibitor); 0.5 μM FCCP (uncoupling agent); and a combination of 0.5 μM rotenone (complex I inhibitor) and 0.5 μM antimycin A (complex III inhibitor). The non-mitochondrial respiratory rate (NMRR) corresponds to the OCR measurement after injection of rotenone and antimycin A. Basal OCR was calculated by subtracting NMRR from the initial recorded OCR (endogenous OCR). Proton leak was calculated by subtracting the NMRR from the OCR following oligomycin injection. Maximal OCR was calculated by subtracting the NMRR from the OCR following FCCP injection.

### Mice

Normal C57BL/6 male mice (7-week-old) were purchased from DooYeol Biotech (Korea), fed a regular chow diet, and housed in a specific pathogen-free animal facility at the temperature of 23 ± 2 °C with a 12 h light/ 12 h dark cycle. For the old mice model, those mice were bred until ∼50-weeks of age. For the HFD model, those normal mice were bred until their weight reached ∼50 g. Sulfasalazine diluted with 1% DMSO was orally administered to the mice at a daily dose of 5 mg/kg (low dose), 50 mg/kg (middle dose), or 500 mg/kg (high dose). Muscle atrophy was induced by immobilization of the hindlimb of normal mice using non-elastic bandage tape (Multipore^TM^ Sports White Athletic Tape, 3M Japan, Japan) and hook-and-loop fastener (Velcro® tape) (one-touch belt, 3M Japan) for 14 days (Velcro model), or by injection of 10 μM cardiotoxin (Sigma-Aldrich) into the tibialis anterior (TA) muscles (CTX model) prior to gavage. The mice were dosed orally with sulfasalazine for 2- or 4-weeks. All animal experiments were conducted in accordance with procedures approved by the Institutional Animal Care and Use Committee of the Chungnam National University.

### Physical performances

Treadmill equipment (Panlab, Harvard Apparatus) was used for treadmill running test. The mice were acclimatized at 15 cm/sec for 10 min at a 10-degree incline for 3 days. After acclimatization, the test was started at 15 cm/sec for 3 min at a 10-degree incline. The speed was accelerated at 5 cm/sec every 3 min to a maximum speed of 40 cm/sec. If mice declined to run, then they were motivated by a transient and mild electric stimulation from the treadmill exercise platform. The total distance traveled by each mouse was calculated at the time the mice became exhausted. The maximal muscle strength of the mice was also determined by measuring grip strength at the appropriate time points using a grip strength meter (Bioseb, Harvard Apparatus). The mice were placed with their hindlimbs or all limbs on a grid and the grip strength was measured immediately before mice fell from the bar. The maximum force was recorded in triplicate. A rotarod apparatus (Panlab, Harvard Apparatus) was used to measure motor coordination and balance. For the accelerating rotarod test (4 - 40 rpm over 300 sec), 3 trials per test were performed during the test day, with a 2 min interval between trials. The latency to fall off the rotarod was recorded for both configurations. Mice that rotated passively were scored as fallen.

### Immunofluorescence and histochemical analysis in tissues

Whole GA muscles were removed from mice that performed physical performance test. For testing of MyHC isoforms in muscle tissue, GA muscle tissues were embedded in O.C.T. compound (Tissue-Tek, SAKURA), frozen in liquid nitrogen-cooled hexane, stored at −80°C, and cut into 10 μm thick cryosections with a cryostat (Leica CM1850, Germany) maintained at −20°C. Immunofluorescence analysis of MyHC isoform expression was performed with primary antibodies against MyHC type I (BA-D5, DSHB), MyHC type IIa (SC-71, DSHB), and MyHC type IIb (BF-F3, DSHB).

Secondary antibodies (Alexa Fluor 488 IgG1 for MyHC IIa, Alexa Fluor 568 IgG2b for MyHC I, and Alexa Fluor 647 IgM for MyHC IIb), which were purchased from Invitrogen, were used for observation with confocal microscope (LSM 880, Carl zeiss, Germany). Percentages of fiber types were calculated by counting muscle fibers positive for each MyHC isoform. For observation the shape of muscle fibers in tissue, muscle tissues were fixed in 4% paraformaldehyde, embedded in paraffin and cut into 6 μm sections. Paraffin-embedded sections were stored at room temperature for later use. Histopathological examinations of hematoxylin-eosin (H&E)-stained GA muscles were performed in paraffin-embedded sections and photographed under a microscope (EVOS^TM^ M5000, Invitrogen). To confirm PAX7 expression in muscle tissue, immunofluorescence was conducted with primary PAX7 antibody (Santa Cruz, sc-81648) and secondary Alexa Fluor 488 antibody (Invitrogen) in paraffin-embedded sections of GA muscle tissue. The nuclei were counterstained with DAPI. Images of all experiments were processed with the Image J software.

## Statistical analysis

Data are presented as the mean ± SEM. Student’s unpaired *t*-tests were performed using GraphPad Prism v.8.0 software. P values < 0.05 were considered statistically significant. Significance is indicated in all figures as follows: * p< 0.05; ** p< 0.01; *** p< 0.001.

## Acknowledgments

This work was supported by the Starting growth Technological R&D Program [TIPS Program, (No. S3198556)] funded by the Ministry of SMEs and Startups (MSS, Korea) in 2021. Jisoo Park was funded by the National Research Foundation of Korea (NRF) grant funded by the Korea Government (MEST) (NRF-2020R1F1A1049801).

The English in this document has been checked by at least two professional editors, both native speakers of English. For a certificate, please see, http://www.textcheck.com/certificate/uKWjKw

## Notes

### Competing Interest Statement

The authors have declared no competing interest.

